# RNase J2 is a specificity factor that governs global mRNA stability in *Bacillus subtilis*

**DOI:** 10.64898/2026.05.27.728192

**Authors:** Ninon Christol, Michelle Goujout, Alexandre Maes, Tania Kamwouo, Hannah LeBlanc, Marie-Françoise Noirot-Gros, Romain Briandet, Ciarán Condon, Sylvain Durand

**Affiliations:** Université Paris Cité, CNRS, Expression Génétique Microbienne (EGM), Institut de Biologie Physico-Chimique, 13 rue Pierre et Marie Curie, 75005 Paris, France; Plateforme de Génomique Fonctionnel, UMR8226 CNRS Sorbonne Université, Institut de Biologie Physico-Chimique, Paris, France; Department of Biology, Massachusetts Institute of Technology, Cambridge, MA 02139.; Université Paris-Saclay, INRAE, AgroParisTech, Micalis Institute, Jouy-en-Josas, France

**Keywords:** RNA degradation, RNase J, *Bacillus subtilis*, Biofilms

## Abstract

Ribonucleases (RNases) play a key role in modulating gene expression at the post-transcriptional level, enabling bacteria to rapidly adapt to their environment. The *B. subtilis* genome encodes more than 20 RNases and understanding the role of each of these enzymes is crucial for a comprehensive knowledge of bacterial post-transcriptional adaptation mechanisms. Among these, RNase J1 and J2, belonging to the ß-lactamase enzyme family, form a heterodimer. While the primary function of RNase J1 as a 5’-exoribonuclease in *B. subtilis* has been known for some time, the role of RNase J2, encoded by the *rnjB* gene, has remained unclear. Indeed, it has been considered a minor degradation factor due its very weak 5’-exoribonuclease *in vitro*, the lack of a growth phenotype and the limited number of mRNAs with altered equilibrium levels in *rnjB* mutants.

In this study, we demonstrate that the absence of RNase J2 influences pellicle and macro-colony biofilm formation, as well as swarming in *B. subtilis*. We show that RNase J2 plays a far more global role in governing mRNA half-lives than originally thought, by stimulating RNase J1-mediated degradation of a subset of RNase J1’s mRNA substrates. An inter-subunit salt-bridge between RNase J1 and J2 is crucial for RNase J2 action. Altogether, our results suggest that RNase J2 acts primarily as a specificity co-factor for RNase J1 rather than as a ribonuclease per se.

**Significance:** This study elucidates the role of RNase J2 in RNA degradation in *B. subtilis*. Through a combination of phenotypic analyses and Rif-seq data, we demonstrate that RNase J2 plays a far more important role than previously appreciated. We provide evidence that physical interactions between RNase J1 and J2 (via salt-bridge) are critical for the degradation of RNase J2 targets, suggesting that RNase J2 acts more as a co-factor of RNase J1 than as a standalone RNase.

## Introduction

Bacteria can adapt to adverse environmental conditions by switching to alternative lifestyles such as biofilm formation, competence development, motility or sporulation. These lifestyle transitions require extensive genetic reprogramming at both transcriptional and post-transcriptional levels, with ribonucleases (RNases) playing a key role in the latter. These enzymes can cleave within transcripts (endoribonucleases) or degrade RNAs from their 5’ or 3’ end (exoribonucleases).

RNases can act on a broad spectrum of mRNA targets, such as the endoribonuclease E in *Escherichia coli*, or the endoribonuclease Y and the exoribonuclease J1 in *Bacillus subtilis*, which play key roles in the RNA degradation process in their respective organisms (1–3). Others have a more limited roles, such as the enzymes involved in ribosomal RNA maturation like RNase M5 or Mini-RNase III in *B. subtilis* (4, 5).

In *B. subtilis*, the main pathway of mRNA degradation initiates with an endoribonucleolytic cleavage by RNase Y, followed by the degradation of the downstream and upstream RNA fragments by the 5’-exoribonuclease RNase J1 and the 3’-exoribonuclease PNPase, respectively. RNase J1 belongs to the ß-CASP sub-family of ß-lactamase enzymes present in all kingdoms of life and has been previously shown to impact more than 30% of the *B. subtilis* transcriptome (1). Intriguingly, most of the ß-lactamase RNases have been shown be capable of both endoribonucleolytic and exoribonucleolytic activities using the same catalytic site (6–9), but RNase J1 is thought to act primarily as a 5’-exoribonuclease *in vivo* in *B. subtilis*.

A paralog of RNase J1, namely RNase J2, is found in the Bacillota and Mollicutes (10). In *B. subtilis*, *Staphylococcus aureus* and *Streptococcus mutans*, RNase J1 has been shown to form a heterodimer with RNase J2 (11–13). Since RNase J2’s 5’-exoribonuclease activity is 1000-fold lower than RNase J1 in *B. subtilis* (11), its exact function within the RNase J1/J2 complex is unclear. Deletion of the *rnjB* gene influences bacterial lifestyle choice such as competence, or biofilm formation in *S. mutans* and *Enterococcus faecalis* (14–16). It also impairs bacterial growth outside of a narrow temperature range around 37°C in *S. aureus* and this has been attributed to a role in stabilizing RNase J1 (16). In contrast, the growth of the *B. subtilis ΔrnjB* strain is unaffected in rich medium at 37°C and a transcriptome analysis showed that the steady state levels of only a dozen mRNAs were up-regulated (17, 18). Interestingly, most of these belonged to the motility Sigma D factor regulon (SigD). *B. subtilis* motility plays a key role in pellicle biofilm development at the air liquid interface (19), but whether the impact of the *ΔrnjB* mutation on the SigD regulon had an effect on biofilm formation was not tested in the original study ^17^.

Here we show that in the absence of RNase J2, pellicle biofilms are thinner, but exhibit a more structured organization. A global RNA half-life analysis revealed an unanticipated impact of the *ΔrnjB* mutation on the stability of thousands of transcripts, including mRNAs encoding proteins important for pellicle formation. Our data suggest that the catalytic activity responsible for the degradation of RNase J2-sensitive RNAs is provided by RNase J1 and that RNase J2 acts as a co-factor for RNase J1 to promote the degradation of a subset of RNase J1 targets in *B. subtilis*.

## Results

### The lack of RNase J2 impacts biofilm formation in B. subtilis

To date, the functional role of RNase J2 in *B. subtilis* remains poorly defined and has long been overlooked, most likely because, in contrast to RNase J1, its absence does not impair bacterial growth under standard conditions in rich medium (2TY) at 37°C (18). However, a previous study in *S. aureus* showed strong growth defects in the *ΔrnjB* mutant strain at 25°C and 42°C (16). To test whether the deletion of RNase J2 had a similar effect in *B. subtilis*, we monitored the growth of WT (non-domesticated strain NCIB3610) and *ΔrnjB* mutant strains in either rich (2TY) or minimal (Spizizen medium), at 25°C or 42°C. (Fig. S1). In contrast to *S. aureus*, only minor effects of the *ΔrnjB* mutation were observed: a ∼40% lower stationary phase plateau in the *ΔrnjB* strain at 42°C in rich medium and about a 25% increase doubling time in minimal Spizizen medium at 25°C (162.1 min *vs* 202.6 min, in WT and *ΔrnjB* strains, respectively).

Prior work also reported that the absence of RNase J2 in *B. subtilis* affects the levels of a limited number of RNAs (17), including the up-regulation of nine genes in the sigma D (SigD) regulon. SigD is a sigma factor involved in the expression of genes related to motility and chemotaxis, two important processes for pellicle biofilm development at the air-liquid interface (20). Moreover, previous studies in *S. mutans* and *E. faecalis* have shown that deletion of RNase J2 impacts biofilm formation (14, 15). Biofilms of *B. subtilis* can formed on agar plates (macrocolonies), at the solid-liquid interface (submerged biofilms) and at the air-liquid interface (pellicles). A distinct genetic program governs the development, architecture, and physiological properties of each type of biofilm (21). To evaluate the impact of RNase J2 on this developmental program in *B. subtilis*, WT and *ΔrnjB* mutant strains were incubated at 37°C in 2TY medium supplemented with glycerol and manganese (2YTGM) to promote pellicle development (22). Under these conditions, the resulting pellicle exhibits a more pronounced wrinkled morphology (Fig. 1A). Confocal imaging of strains constitutively expressing GFP from the *amyE* locus and grown in 2YTGM medium at 30°C, enabled detailed visualization of pellicle formation kinetics. The WT strain reached the air-liquid interface slightly earlier than the *ΔrnjB* strain (11h *vs* 12h of pellicle development). However, after 14h, the *ΔrnjB* strain exhibited an increased density of fluorescence at the air-liquid interface (Fig.1B) and quantification of pellicle roughness confirmed that the *ΔrnjB* strain formed a more wrinkled pellicle (Fig. 1C). In contrast, the WT strain displayed a more diffuse fluorescence signal, consistent with the formation of a thicker pellicle, as evidenced by its greater biovolume (Fig. 1C). Altogether, these results suggest that a lack of RNase J2 increases the number of cells migrating to the air-liquid interface, contributing to the enhanced wrinkling observed in the pellicle. These observations align with the previously reported up-regulation of the SigD regulon in the *ΔrnjB* mutant strain.

**Figure 1.**
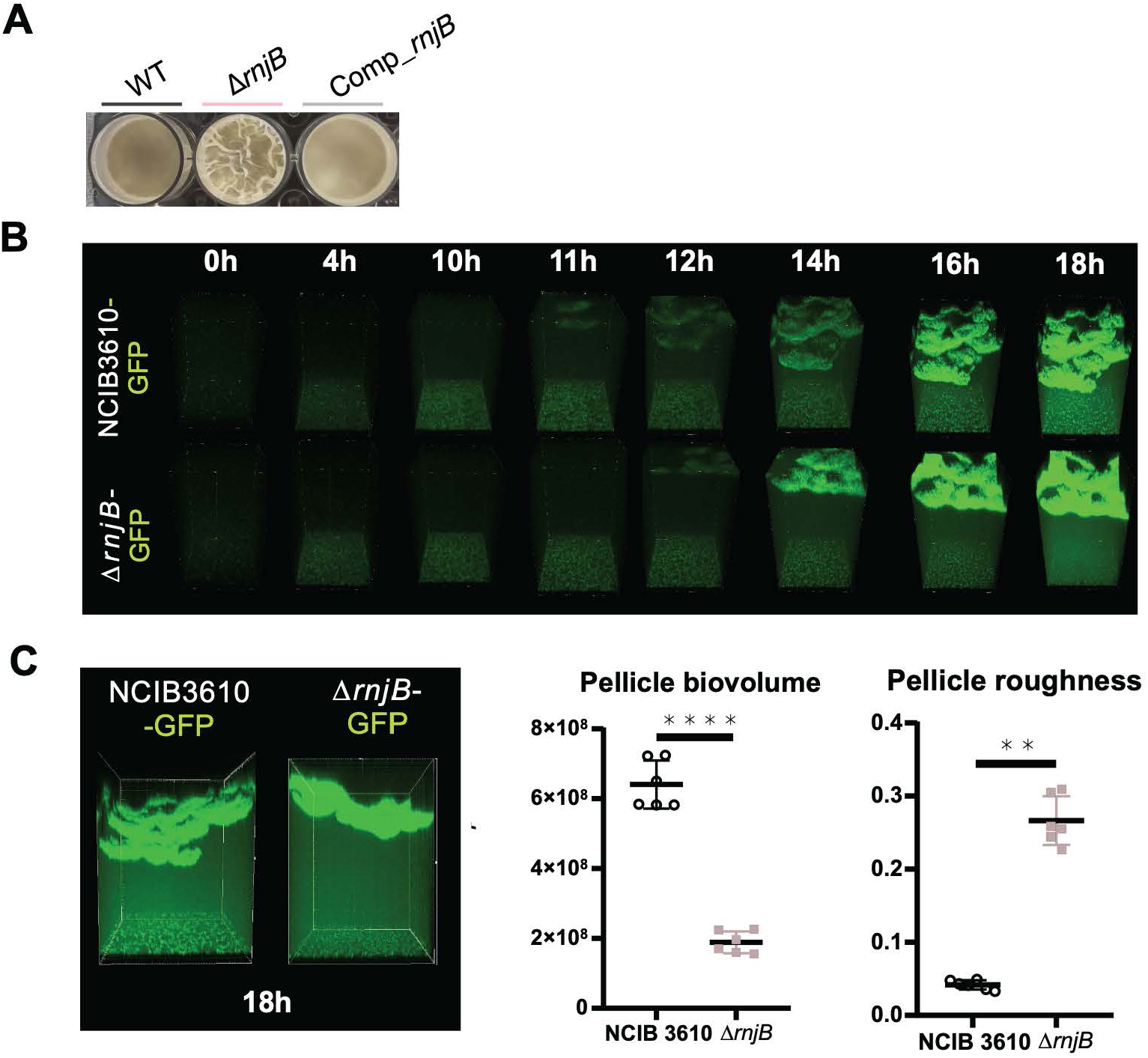
Impact of rnjB deletion on pellicle formation. (A) NCIB3610 wild-type (WT), *ΔrnjB* and *ΔrnjB* complemented strains (amyE::rnjB) were grown in rich medium (2YT) to mid-exponential phase (OD_600_ 0.6) before being diluted to OD_600_ 0.01 in 2YTGM [2YT complemented with 0.1 mM MnSO4 and 1% glycerol] in 24-wells plates and incubated at 37°C for 24h. (B) Spatial confocal imaging was performed on pellicles formed from 0 to 18 hours at 30°C in 2YTGM, using NCIB3610 wild-type (WT) or *ΔrnjB* strains genetically modified to constitutively express the GFP protein (green). (C) Spatial confocal microscopy acquisitions were used to quantify the biovolume and roughness of pellicles after 18 hours of development. At least three independent biological replicates were performed for each condition, with two independent microscopic fields acquired per biofilm well. All quantifications were conducted on the complete dataset. A representative image was selected and is shown in the figure.

We also recorded the development of macro-colony biofilms and compared macro-colony area (N) and growth rate (µ) in the WT and the *ΔrnjB* strains (Fig. 2A, B). The *ΔrnjB* mutant strain grew faster on average (µmax=0.19 mm^2^/h vs 0.15 mm^2^/h) and occupied a significantly larger final area (Nmax=248 mm^2^ vs 183 mm^2^) than the WT strain (Fig. 2A). The growth dynamics also differed markedly between the two strains; the WT strain exhibited biphasic growth, whereas the *ΔrnjB* strain displayed a single, prolonged and more pronounced growth phase (Fig. 2B).

**Figure 2.**
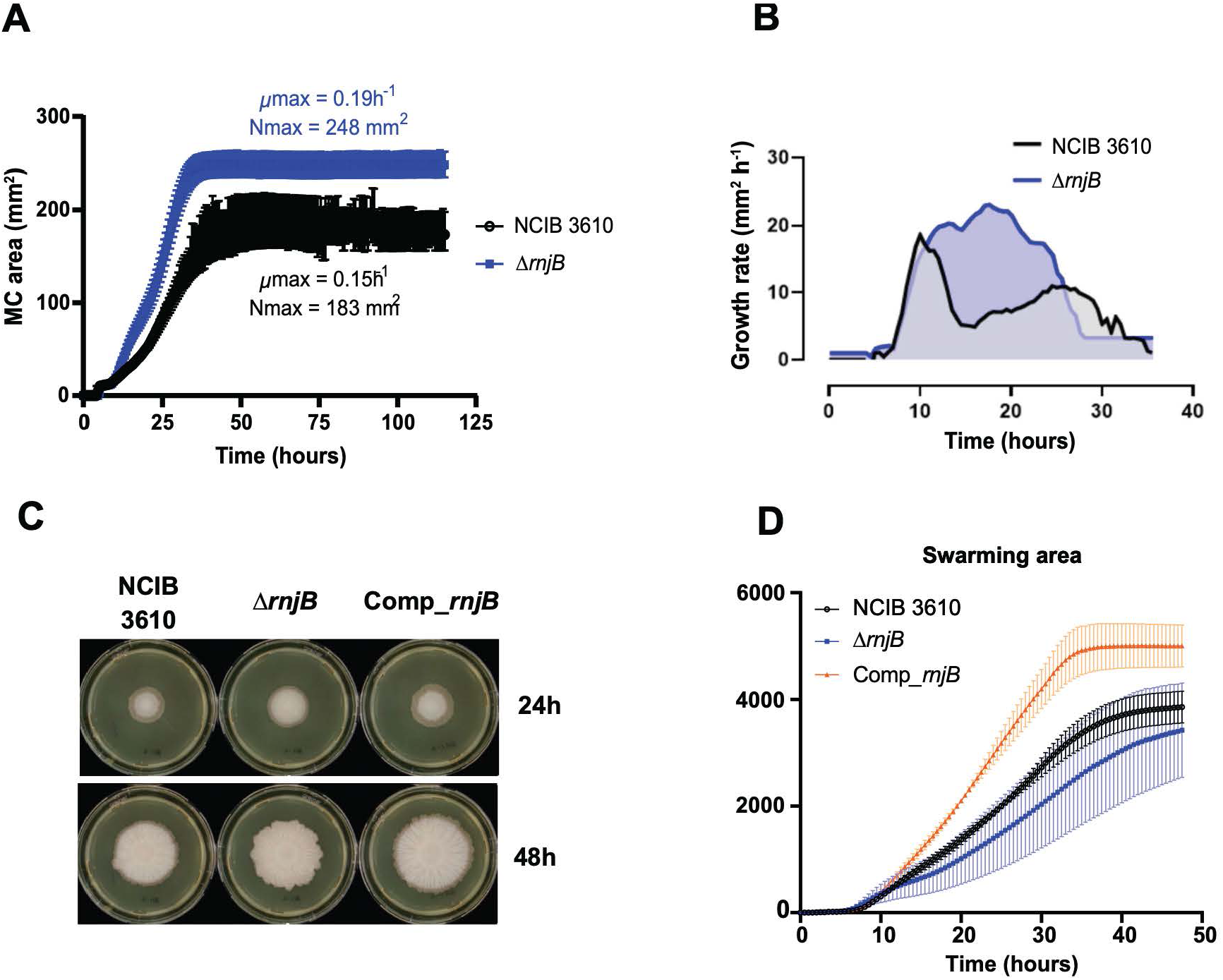
Macro-colony development and swarming over time. (**A**) Development of the microcolony area and **(B)** analysis of macro-colony propagation velocity (μ) of the wild-type and ΔrnjB strains over time over time. (**C**) Screenshot of the swarming video at 24h and 48 h of NCIB3610, *ΔrnjB* and *ΔrnjB* complemented strains in 2XYTGM at 37°C (**D**) Quantification of swarming area over time.

Given that both pellicle and macro-colony development were modified in the *ΔrnjB* strain compared to WT, we also investigated the swarming capacity of these two strains. Although the swarming area was statistically comparable between the WT and the *ΔrnjB* strains, the cell density at the periphery of the swarming area was consistently higher in the *ΔrnjB* strain throughout migration (Fig. 2C). These findings suggest that the absence of RNase J2 does not enhance individual cell motility but rather increases the proportion of cells adopting a motility lifestyle compared to the WT strain. This interpretation aligns with the higher cell concentration at the air-liquid pellicle interface observed in the *ΔrnjB* strain as motility is a critical factor for cell migration in these conditions. Interestingly, the strain complemented for RNase J2 presents an increased swarming area compared to the WT strain (Fig. 2C, D), suggesting enhanced individual cell motility more than an altered proportion of motile cells in the population, given that cell density at the periphery of the swarming area remains comparable to that of the WT strain. This difference correlates to the increased expression of RNase J2 in the complemented strain compared to the WT strain (Fig. S2).

### RNase J2 is a key post-transcriptional regulator with a broad spectrum of RNA substrates

The previous global transcriptome analysis comparing WT and *ΔrnjB* strains showed only 44 transcripts with altered abundance in the absence of RNase J2 (32 down– and 12 up-regulated) (17). However, this number is likely to be an underestimate, since the study was conducted using the domesticated *B. subtilis* 168 strain, which is known to be non-motile and to exhibit a reduced ability to form architecturally complex and robust biofilms compared to non-domesticated strains such as NCIB3610. Secondly, the study analyzed steady-state RNA levels during exponential growth in LB medium. However, a recent study in *Salmonella* demonstrated that steady-state RNA abundance can be largely uncorrelated with changes in RNA half-life in strains lacking specific RNA-binding proteins (23).

To gain a deeper insight into the role of RNase J2 in the RNA degradation in *B. subtilis*, we performed a Rif-seq experiment to directly measure RNAs half-lives in both WT and *ΔrnjB* strains in 2TY medium, following an approach similar to that applied in *Salmonella*. Total RNA was extracted at time 0 (just prior to rifampicin addition) and 1, 2, 3, 5 and 10 minutes after rifampicin treatment, to inhibit *de novo* transcription. Half-lives were calculated from semi-log plots of the percent RNA remaining over time for each gene or non-coding RNA segment, annotated based on NCIB3610 reference genome (GCF_002055965.1) completed with RNA “segments” and “regulatory RNA” features from strain 168 (SubtiWiki, GCF_000009045.1)(Fig. S3). The median half-life was around 1.7 min in the WT strain, less than the 5 min previously determined by DNA microarrays (24), but slightly higher than was observed recently in *Salmonella* (0.9 min) with a similar Rif-seq study (23). In the absence of RNase J2, the median half-life increased to 3.3 min (Fig. 3). Strikingly, RNase J2 deletion increased the half-life of 1,911 RNAs (30% of all annotated features) by more than 1.5-fold compared to the wild-type strain (Fig. 4 and Table S1). All categories of RNAs (genes and non-coding RNAs) were affected by RNase J2 deletion.

**Figure 3.**
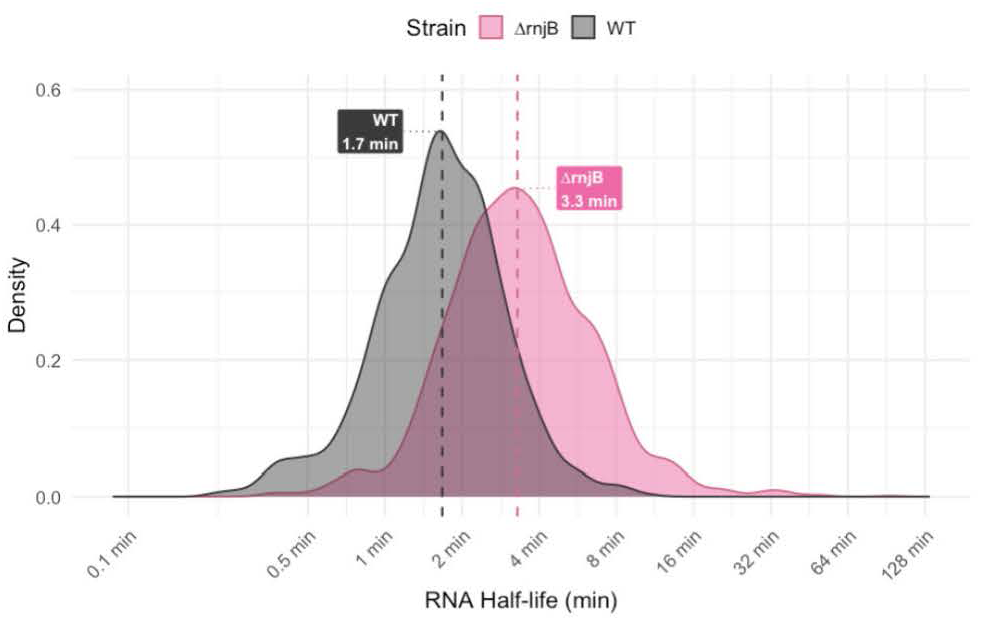
Half-life distribution in WT and ΔrnjB strain. Density plot comparing the distribution of mRNA half-lives (HL) between the WT (black) and *ΔrnjB* strains (pink). Only the mRNA half-life with a mean absolute error ≤0,6 are represented. The median mRNA half-life for each strain is indicated above the respective curve.

**Figure 4.**
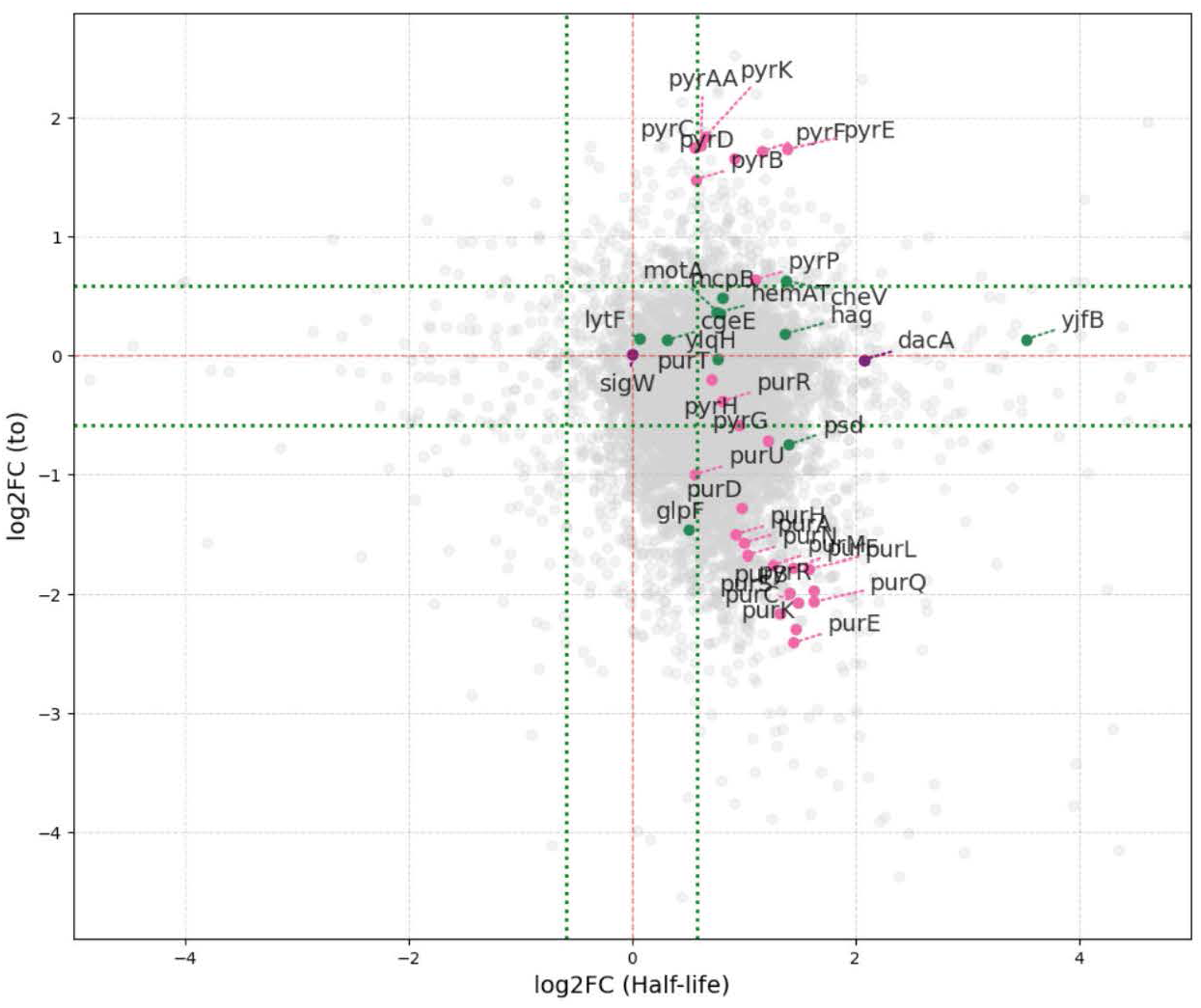
Scatter plot comparing log2 fold-changes (log₂FC) between gene expression determined at T0 and mRNA half-life between the wild-type (WT) and *ΔrnjB* mutant strains of *B. subtilis*. The green dashed line represents a +/− 1.5-fold change cutoff (log2FC= +/−0,585). Genes from the pur and pyr operons are highlighted in pink, those previously identified as upregulated in study ^17^ are shown in green, and additional genes (dacA, sigW) further analysed in the study are indicated in violet.

Among the nine SigD-dependent genes upregulated in the absence of RNase J2 identified by Mäder *et al* (17), five exhibited a change in half-life greater than 1.5-fold (p-value ≤0.05) in our Rif-seq. analysis (Fig. 4). This was the case for *motA* (1.7-fold stabilization), for example, encoding a part of the flagellar stator and required for efficient pellicle biofilm formation (20) and two genes linked to chemotaxis and involved in biofilm formation: the *hemAT* transcript encoding a soluble chemotaxis receptor that senses oxygen (20) and the *cheV* transcript, encoding a modulator of CheA activity in response to attractants (1.7-fold and 2.6-fold, respectively). Our Rif-seq analysis also identified other mRNA involved in biofilm formation with increased half-lives: the *degU* mRNA, encoding a two-component response regulator (25) (1.7-fold), and some genes of the *eps* operon, encoding enzymes involved in extracellular polysaccharide synthesis (26) (up to 3.3-fold stabilization). The stabilization of these mRNAs in the absence of RNase J2 is consistent with the enhanced localization of *B. subtilis* cells to the air-liquid interface and the increased wrinkling of the pellicle.

The lack of RNase J2 also led to an increased stability of several mRNAs encoding key transcriptional regulators. These include the *rpoD* mRNA, encoding the primary sigma factor SigA, and the *sigE* and *sigG* mRNAs, encoding sporulation-specific sigma factors (2-fold, 1.9-fold, and 1.8-fold, respectively). We did not see a reproducible effect on the half-life of the *sigD* mRNA, the second last cistron of a 20 kb operon. However, the stability of the mRNA segments encoding eight of the first ten cistrons of this operon *fliE*-*fliF*-*fliG*-*fliH*-*fliI*-*fliJ*-*ylxF*-*fliK*, was increased between 1.6-fold and 3.5-fold. The gradual increase in the half-life of promoter-distal genes within very long operons is attributed to ongoing transcription that is not inhibited by rifampicin, thereby artificially inflating the estimated half-life values. The mRNAs of other important transcriptional regulators were also stabilized in the *ΔrnjB* mutant strain, such as CodY, a pleiotropic transcriptional repressor that senses limitations in both GTP levels and branched-chain amino acids (2.6-fold); ResDE a two-component system that regulates aerobic and anaerobic respiration (2-fold) and PurR, a transcriptional repressor that regulates purine biosynthesis (1.7-fold).

As expected for the lack of an RNase, only 3 mRNAs showed a >1.5-fold decrease in half-life in the absence of RNase J2, suggesting that its primary role is to participate in mRNA degradation.

Since RNase J2 is involved in the degradation of mRNAs encoding several transcriptional regulators (see above), we compared changes in steady state RNAs levels at T0 to changes in RNA half-lives, to distinguish transcriptional from post-transcriptional effects (Fig. 4). Of a total of 1,363 RNAs whose equilibrium levels were significantly altered at T0 in the *ΔrnjB* strain, 1,229 (90%) were reduced; yet only 3 of these RNAs showed decreased stability. Indeed, only 57 RNAs showed a coherent change (both increased or both decreased) in steady-state levels (T0) and half-life (|log₂FC| ≥ 0.585, p-value ≤ 0.05, MAE ≤ 0.6). These results indicate a substantial decrease in global transcription in the absence of RNase J2. Interestingly, 78% of tRNAs were downregulated at steady-state (T0) (Fig. S4).

In many cases (485), the transcriptional down-effect counteracted a stabilization up-effect caused by the loss of RNase J2. This effect was exemplified in the set of genes regulated by the transcriptional repressor PurR. Indeed, 21 of the 28 mRNAs of the PurR regulon, including the different coding segments of the long *purE-K-B-C-S-Q-L-F-M-N-H-D* operon, had increased mRNA stability that was counteracted by a stronger decrease in equilibrium transcription levels at T0. (Fig. 4 and S5). One exception is the *pdxST* operon, encoding a pyridoxal-5-phosphate synthase, where no transcriptional effect was observed. This may be linked to stabilizing effect of the *ΔrnjB* deletion on the mRNA coming from the upstream gene *dacA* (see below).

In contrast to the negative impact of the *ΔrnjB* deletion on the transcription of genes involved in purine biosynthesis, mRNAs involved in the pyrimidine-synthesis pathway were up-regulated in the *ΔrnjB* strain at T0. The downregulation of the transcription repressor PyrR (3.9-fold) at T0 likely explains the increased expression of the *pyr* operon (Fig. 4 and S5). Interestingly, a similar increased in expression of the *pyr* operon was observed in the absence of RNase J2 in *E. faecalis* (15), suggesting that this function of RNase J2 is conserved in Gram-positive bacteria.

### Validation of Rif-seq analysis by Northern blotting

To validate our Rif-seq analysis, we compared RNA half-lives estimated from the transcriptome data with those measured by Northern blot for a subset of selected transcripts. Among the transcripts analyzed were *dacA*, which encodes a major D-alanyl-D-alanine carboxypeptidase and was among the 30 most stabilized RNAs in our Rif-seq dataset. We also included *motAB,* for its involvement in motility and biofilm formation (27), and *cheV,* since it is involved in chemotaxis and plays a role in pellicle biofilm formation (19). In addition, we also validated the impact of RNase J2 on the half-life of the mRNA encoding the PurR repressor. The RNA half-lives determined by Rif-seq and Northern blot for the selected transcripts are presented Fig. 5 and S6.

**Figure 5.**
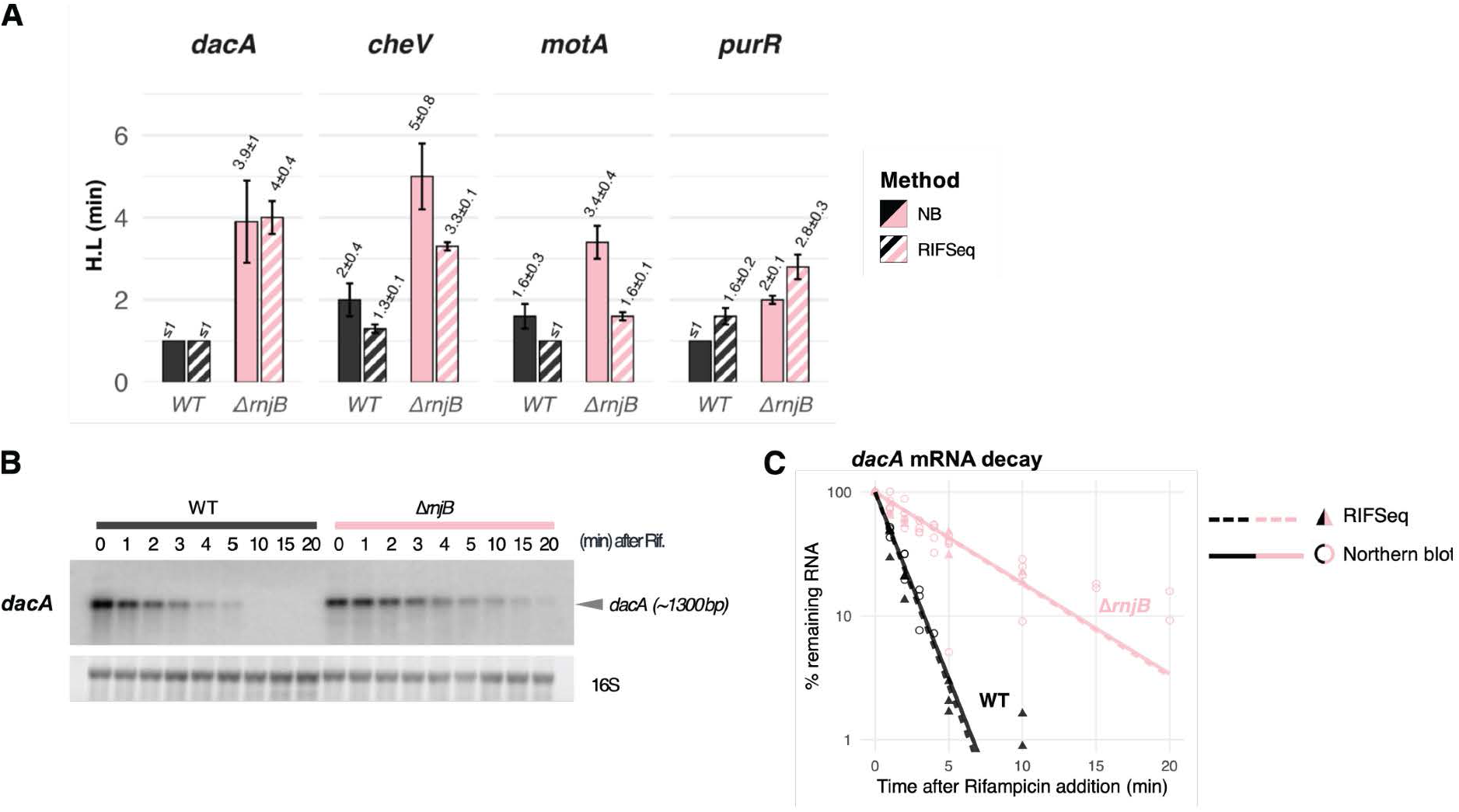
Comparison of mRNA half-lives (HL, in minutes) estimated from Northern blots (NB) and from the RIF-Seq data for selected transcripts. **(A)** mRNA stabilities of *dacA*, *cheV*, *motA* and *purR* mRNAs in *B. subtilis* strains WT (black) and *ΔrnjB* strains (pink). Solid bars represent NB-derived half-lives; hatched bars represent RIF-Seq-derived half-lives. Error bars indicate ± standard deviation (SD). **(B)** Representative Northern blot probed with a *dacA*-specific probe. **(C)** Quantification of signals from Northern blots probed with a *dacA*-specific probe, overlaid with the corresponding RIF-Seq decay curves. Individual biological replicates are represented as transparent open circles for Northern blot data and as triangles for RIF-seq experiments; the lines represent the exponential decay of the dacA mRNA in both strains.

The half-lives calculated by Rif-seq were quite similar for *dacA* and *purR* mRNA to those measured by Northern blot analysis and the half-lives for *motA* and *cheV* in the WT and *ΔrnjB* strains derived from Rif-seq data tended to be lower. However, the relative trends in RNA stability were consistently preserved in both approaches. These findings confirm the robustness of our Rif-seq experiment. Taken together, our results indicate that, although deletion of *rnjB* does not impair growth in rich medium, it plays a far more extensive global function in RNA turnover than previously thought.

The similar levels of the *pdxST* mRNA in WT and *ΔrnjB* strains (see above) could be due to the stabilization of the *dacA* mRNA. Indeed, transcription of *pdxST* is driven by two promoters: a proximal promoter located immediately upstream of the *pdxS* gene and regulated by PurR, and a more distal promoter upstream of *dacA*, which produces a longer transcript encompassing the *dacA-pdxS-pdxT-serS* coding sequences. Consequently, the increased stability of the *dacA*-derived transcript may compensate for PurR-mediated transcriptional down-regulation of the downstream *pdxST* genes.

### The predicted catalytic site of RNase J2 is not necessary for degradation of its targets

All categories of RNAs (genes and non-coding RNAs) were affected by RNase J2 deletion. Given the involvement of RNase J2 in the degradation of about 1/3 of *B. subtilis* RNAs (1,911 / 6,170 annotated RNAs), we sought to clarify its specific function in mRNA turnover by asking whether its predicted catalytic site directly contributes to mRNA degradation. The signature zinc-binding motif of the ß-lactamase catalytic site (HxHxDH) is partially degenerate in RNase J2 (^74^HGHDEN^79^) compared to RNase J1 (^74^HGHEDH^79^). *In vitro* assays have shown that RNase J2 exhibits 1000-fold weaker 5’-exoribonuclease activity than RNase J1 and that both enzymes have a very weak endoribonucleolytic activity, which can be abolished by the H76A point mutation (28, 29). To investigate the role of the RNase J2 catalytic site, we measured by Northern blot the half-lives of the *dacA*, and *cheV* mRNAs in a strain expressing the H76A catalytic-site mutant as the only source of RNase J2. Surprisingly, the half-lives of these two transcripts remained completely unchanged in the strain carrying the RNase J2 H76A mutation compared to the WT strain (Fig. 6). These observations suggest that RNase J2 is crucial for the efficient degradation of these targets, not through its catalytic activity, but rather through its physical presence.

**Figure 6.**
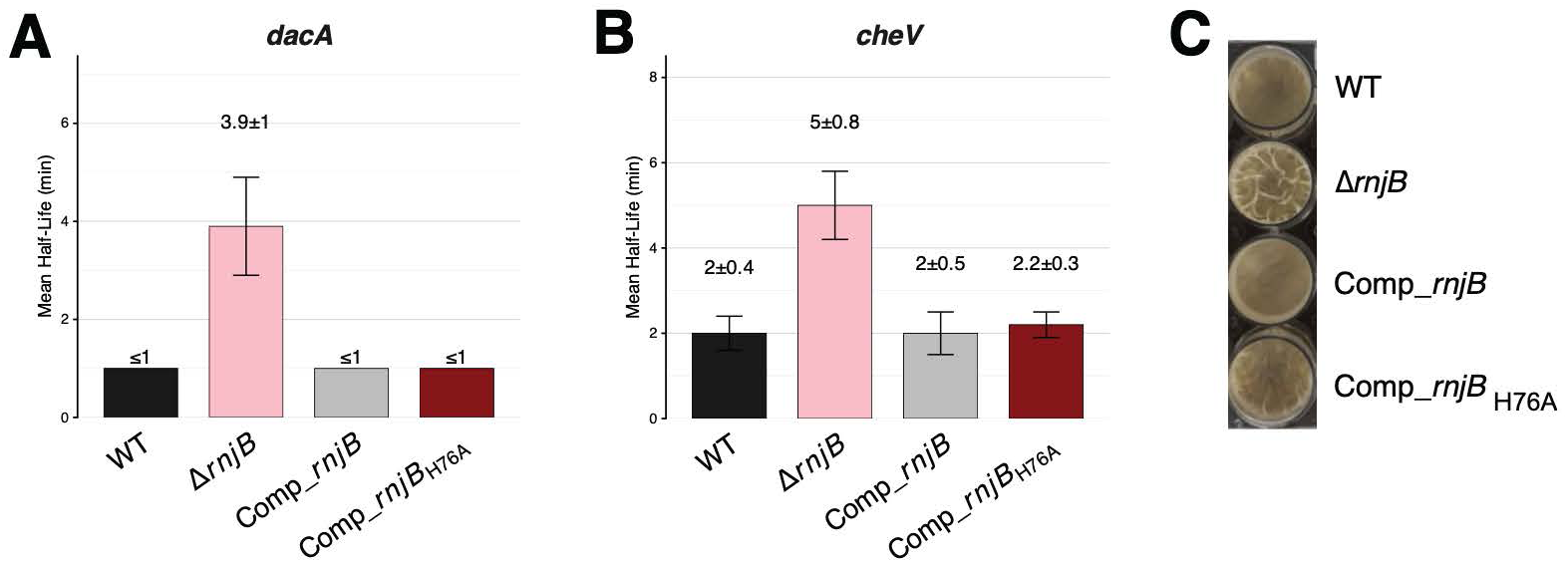
Impact of the RNase J2 catalytic-site mutants on mRNA half-lives and pellicle formation. (**A, B**) Histograms comparing the half-lives of *dacA* (**A**) and *cheV* (**B**) mRNAs in WT, *ΔrnjB* and *ΔrnjB* strains complemented with either WT *rnjB* or the catalytic-site mutant *rnjB* (H76A) expressed from the *amyE* locus. Samples were collected at mid-exponential phase in 2YT medium, before (0) and after rifampicin treatment (1 to 20 min time points). Half-lives were quantified from Northern blot membranes probed for *dacA* and cheV. Representative Northern blots for each quantification are presented Fig. S5. Half-lives were calculated from at least two independant experiments. (**C**) B. subtilis NCIB3610 wild-type (WT), *ΔrnjB* and *ΔrnjB* strains complemented with either WT *rnjB* or *rnjB* (H76A) were grown in rich medium (2YT) until mid-exponential phase (OD_600_ 0.6) before being diluted to OD_600_ 0.01 in 2YTGM [2YT supplemented with 0.1 mM MnSO4 and 1% glycerol] in 24-well plates. Plates were sealed with parafilm, to avoid oxygen concentration differences between wells, and incubated at 37°C for 24h. One representative image of three biological replicates is shown.

We also evaluated the impact of the RNase J2 catalytic-site mutant on pellicle formation. Interestingly, this mutant displayed an intermediary phenotype, with wrinkles slightly more visible than the WT but not as much as in the *ΔrnjB* mutant strain (Fig. 6C).

### RNase J2 promotes RNase J1 degradation of a subset of its mRNA substrates

Since RNase J2 forms a heterodimer with RNase J1, we investigated how deletion of the RNase J1 gene (*rnjA*) affected the half-lives of selected transcripts (*dacA*, *purR* and *cheV*) whose stability was impacted by the *ΔrnjB* deletion (Fig. 7A). The half-lives of the *dacA* and *cheV* mRNAs were similarly increased in the *ΔrnjA*, *ΔrnjB* and *ΔrnjA ΔrnjB* double mutant strains (∼4 min for *dacA* and ∼5-6 min for *cheV*) compared to WT (≤1min for *dacA* and 2 min for *cheV*). This suggests that RNase J2 is fully necessary for the degradation of the *dacA* and *cheV* mRNAs by RNase J1. The *purR* mRNA, on the other hand, exhibited a markedly longer half-life in the Δ*rnjA and ΔrnjA ΔrnjB* double mutant (∼8-10 min) compared to the *ΔrnjB* mutant strain (2 min). Nevertheless, all three mutants showed increased stability of the *purR* mRNA relative to the WT strain (≤1 min). These results suggest that while RNase J2 is necessary for the efficient degradation of some RNase J1 substrates, the degradation of other transcripts, such as *purR,* can occur, at least in part, independently of RNase J2.

**Figure 7.**
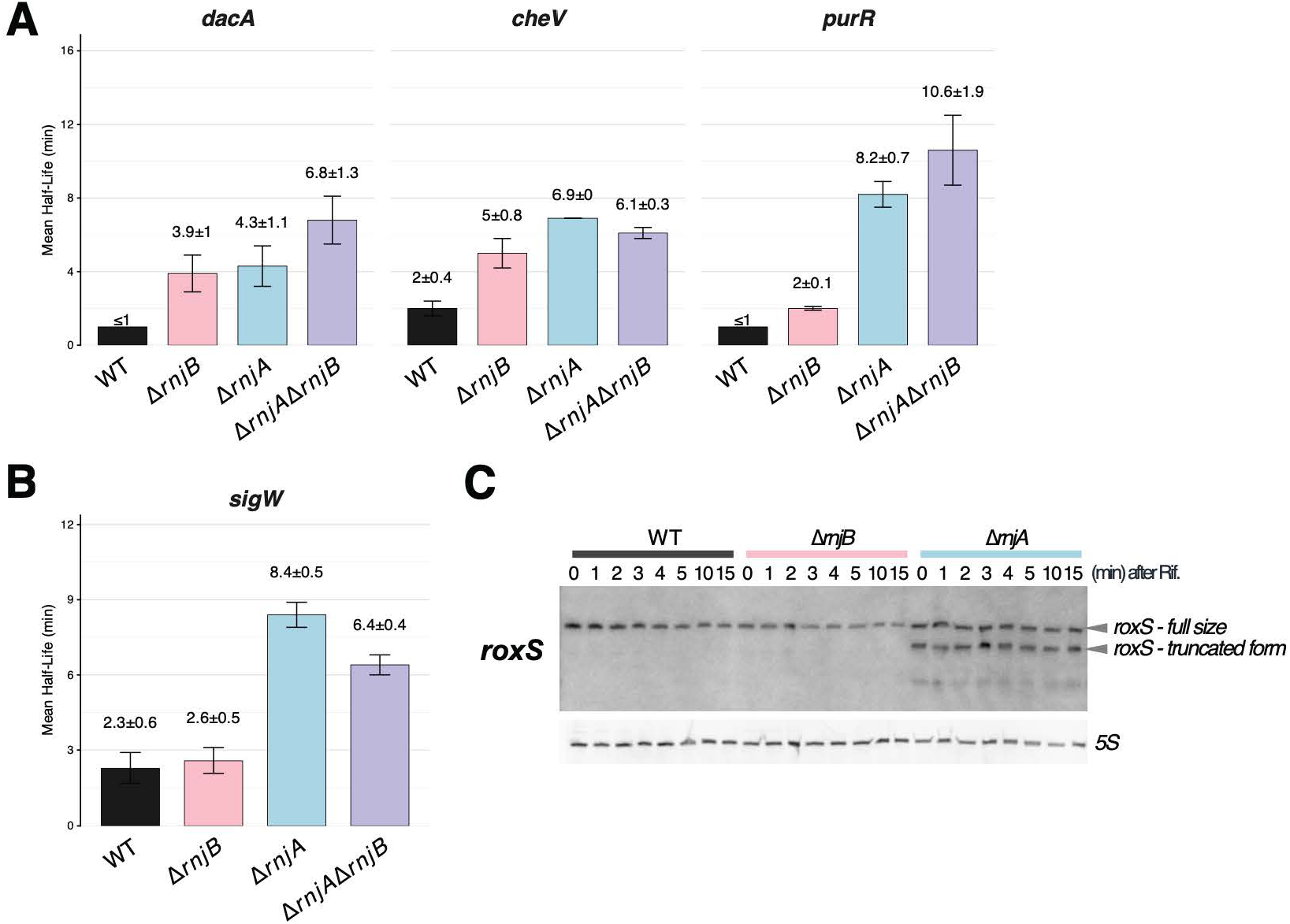
RNase J2 promotes RNase J1 degradation of a subset of its mRNA substrates. (**A**) Effect of RNase J1 on RNase J2 targets identified by RifSeq. Half-lives (in minutes) of *dacA*, *purR*, *cheV*, RNAs in *B. subtilis* Wild-Type (WT), *ΔrnjA*, *ΔrnjB* and *ΔrnjA/ΔrnjB* double mutant strains, calculated from Northern blots probed with radiolabelled oligonucleotides. Error bars indicate ± SD. **(B)** and **(C)** Effect of RNase J1 on non-RNase J2 targets. (**B**) Comparison of mRNA half-lives (H.L, in minutes) estimated from Northern blots for the *sigW* transcript. Error bars indicate ± SD. **(C)** Northern blot of total RNA from WT, *ΔrnjB* and *ΔrnjA* strains, probed with a roxS radiolabelled probe. All samples for Northern blots were collected at mid-exponential phase in 2YT medium, before (0) and after rifampicin treatment (1 to 20 min time points).

To determine whether all targets of RNase J1 require RNase J2 as a co-factor as a general rule, we measured the RNase J2-dependence of the half-lives of two RNase J1 targets identified in previous studies (1, 30). These were the *sigW* mRNA, encoding a sigma factor required for the adaptation to membrane stress and RoxS, a non-coding RNA involved in regulating NAD+/NADH ratios in *B. subtilis*. Neither RNA was identified as an RNase J2 target in our Rif-seq dataset (Fig.4 and Table S1). Northern blot analysis confirmed that stability of the *sigW* RNA was stabilized in the absence of RNase J1, but completely insensitive to RNase J2 (∼2 min half-life in WT and *ΔrnjB* strains vs ∼6-8 min in *ΔrnjA* and *ΔrnjA ΔrnjB* double mutants) (Fig. 7B). Similarly, the two main RNase J1-dependent degradation intermediates of RoxS accumulated in the *ΔrnjA* strain, as seen previously, but not in the *ΔrnjB* deletion mutant (Fig. 7C). Together, these results suggest that RNase J2 stimulates RNase J1 degradation of only a subset of RNase J1’s targets.

### Efficient degradation of RNase J2 targets requires functional contact with RNase J1

To confirm that RNase J2 stimulates RNase J1 activity through direct interaction between the two proteins, we targeted an arginine residue at position 544 and aspartic acid 449 of RNase J2, since these two residues have been shown to form salt bridges at the homodimer interface in the crystal structure of *B. subtilis* RNase J1 (31) (Fig. 8A). An AlphaFold model suggests that these inter-subunit salt-bridges are maintained in the RNase J1/J2 heterodimer. We therefore mutated arginine 544 to glutamic acid (R544E) and aspartic acid 449 to a lysine (D449K) in the complementing copy of *rnjB* in the *amyE* locus.

**Figure 8.**
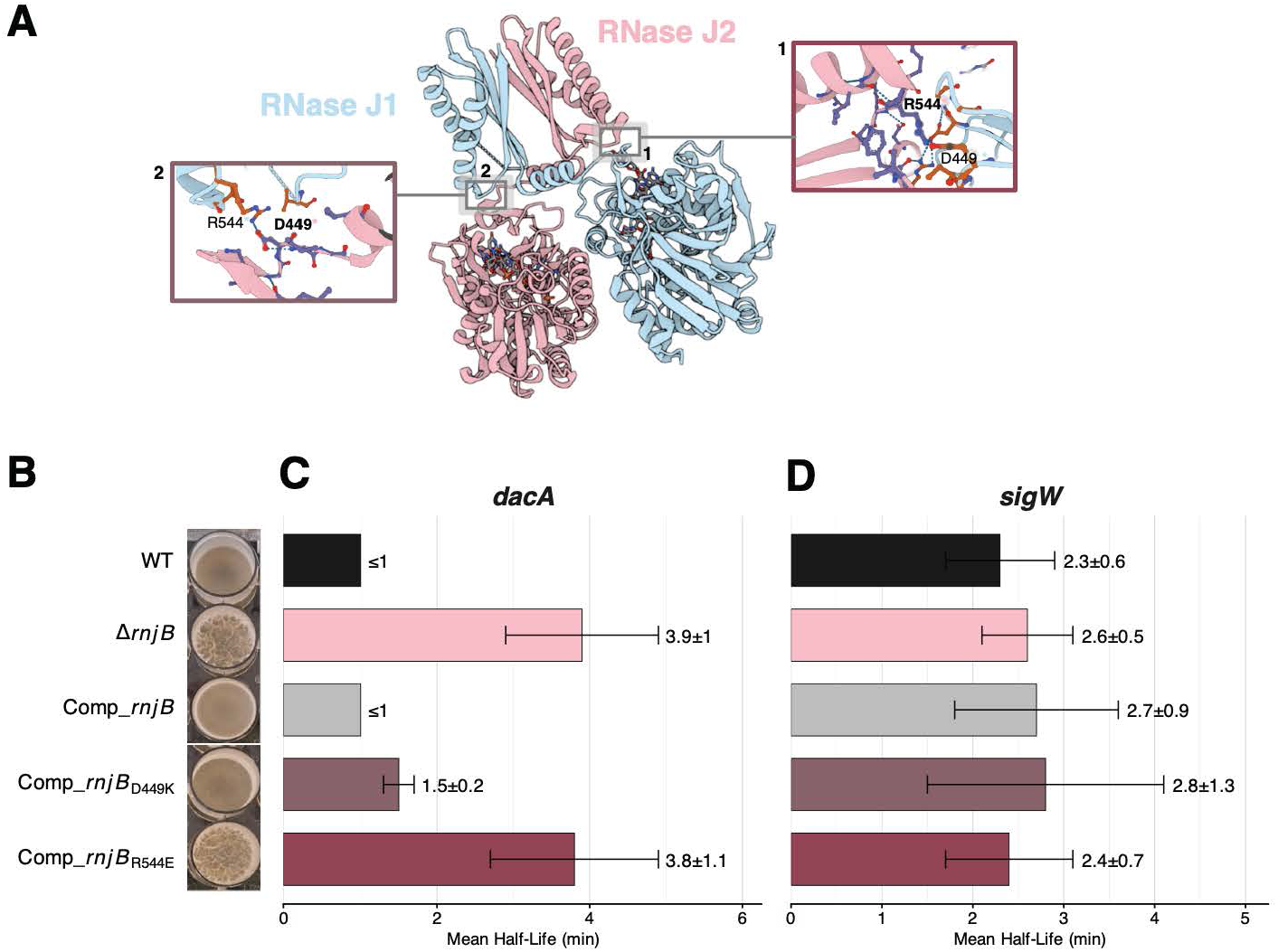
Impact of mutations in RNase J2 residues involved in interaction with RNase J1 on pellicle formation and mRNA stability. (**A**) Visualisation of interaction sites between RNase J1 and RNase J2. The predicted Alphafold structure of RNase J2 was assembled to the experimental structure of RNase J1 (3ZQ4) on Pymol and visualized with Mol*Viewer. RNase J2 residues are shown in bold and pink *, RNase J1 in blue regular font marked *.(**B**) WT, *ΔrnjB*, and *ΔrnjB* strains complemented with RNase WT *rnjB* or *rnjB* interaction mutants were grown in rich medium (2YT) to mid-exponential phase (OD_600_ 0.6) before being diluted to OD_600_ 0.01 in 2YTGM [2YT supplemented with 0.1 mM MnSO4 and 1% glycerol] in 24 wells plates and incubated at 37°C. (**C** and **D**) Comparison of mRNA half-lives estimated from Northern blots of total RNA extracted from WT, *ΔrnjB* and *ΔrnjB* strains complemented with genes expressing C-terminal His tagged version of WT RNase J2, RNase J2 (D449K) or RNase J2 (R544E) at the amyE locus. Samples were collected at mid-exponential phase in 2YT medium, before (0) and after rifampicin treatment (1 to 20 min time points). The membranes were probed for *dacA* and *sigW* with a radiolabelled probe. Half-life calculation for *dacA* (**C**) and *sigW* (**D**) is indicated in the histogram. A representative membrane is shown Fig. S5.

We first asked whether these mutations affected the stability of the *rnjB* transcript or the cellular level of the RNase J2 protein (Fig. S2). Neither mutant affected *rnjB* mRNA stability, and although the R544E-complemented mutant showed a modest decrease in protein abundance compared to either the WT– or D449K-complemented strain, its protein level remained higher than when expressed from the native locus. We next tested how these mutations affected pellicle biofilm formation (see Fig. 8B). The structure and pellicle formation kinetics of the D449K mutant closely resembled those of the WT strain, whereas the R544E mutant displayed pellicle characteristics similar to the *ΔrnjB* strain. To confirm the impact of the R544E mutation on RNase J2 mRNA target degradation, we first measured the half-life of the *dacA* mRNA targets in this mutant (Fig.8C, D). A similar half-life was calculated for the *dacA* mRNA in both the *ΔrnjB* and R544E mutant strains (∼4 min), which was more stable than that of the WT strain (≤1 min) or the D449K mutant strain (∼1.5 min). To further investigate the specificity of this interaction, we also examined the impact of these two mutants on the stability of *sigW*, a known RNase J1-specific target (see above). Remarkably, neither mutant had a significant impact on *sigW* mRNA stability. These results confirmed that the inter-enzyme contact between RNase J1 and RNase J2 via R544E is essential for the degradation of RNase J2 targets by RNase J1, but is dispensable for substrates exclusively degraded by RNase J1.

## Discussion

Until now, the role of RNase J2 in *B. subtilis* has been considered accessory, as its deletion has not been associated with any obvious growth phenotype and its 5’-exoribonucleolytic activity *in vitro* was three orders of magnitude lower than that of RNase J1. Moreover, only a handful of potential targets were identified in a previous transcriptome analysis. The present study uncovered an unexpected yet important role for RNase J2 in RNA degradation in *B. subtilis*. Using Rif-seq analysis, we show that the lack of RNase J2 alters the stability of over a thousand mRNAs *in vivo*. Measuring RNA half-lives after transcriptional arrest with rifampicin has already proven to be a powerful technique to uncover post-transcriptional effects, as shown in *Salmonella* (23). This previous study demonstrated that RNA-binding proteins such as ProQ have a much greater impact on RNA stability than is observed under steady state conditions. Indeed, changes in RNA half-lives often result in little to no effect on RNA abundance at steady state. The reason for this observation in bacteria is still unclear and may result from compensatory mechanisms, such as a decrease in transcription following RNase J2 deletion, as suggested by the down-regulation of thousands of genes at T0, including 78% of tRNAs (Fig. S5). This phenomenon has been described as transcriptional buffering in yeast (32). Indeed, it was shown that deletion of several degradation factors, including the 5’ exoribonuclease Xrn1, inhibits transcription initiation and elongation (33, 34).

Deletion of RNase J2 impacts the kinetics and architecture of pellicle biofilm at the air-liquid interface, as well as macro-colony development. Interestingly, the swarming ability of individual cells appears to increase in parallel with RNase J2 expression in the complemented strain (Fig. 2C). One possible explanation for this observation is the enhanced degradation of *slrR* mRNA in the complemented strain, as *slrR* encodes an important negative regulator of flagellar gene expression and has been identified as a target of RNase J2 (Table S1). Taken together, these results echo findings in other bacteria where loss of RNase J2 affects cell fate, such as biofilm formation. Furthermore, our work shows that deletion of RNase J2 strongly up-regulates the pyrimidine-biosynthesis pathway, mirroring observations in *E. faecalis* (15). However, in this bacterium, it was suggested that this effect is mediated by the transcriptional repressor PyrR. Surprisingly, the authors reported an up-regulation of *pyrR* mRNA levels by qRT-PCR. In contrast, our experiments revealed a decrease in *pyrR* mRNA levels in the *ΔrnjB* mutant, more consistent with the observed up-regulation of the *pyrR* operon.

Although the PurR mRNA is stabilized in the *ΔrnjB* strain, its steady-state level remains unchanged, likely due to autoregulation that compensates for the increased stability. This suggests that the significant transcriptional impact of RNase J2 observed on the *purR* regulon at T0 may result from alterations in the level of PurR’s effector rather than changes in PurR itself. Indeed, PurR DNA-binding is inhibited by phosphoribosylpyrophosphate (PRPP). Given that the level of the phosphoribosylpyrophosphate synthase (*prs*) is reduced 1.9-fold at steady state in the absence of RNase J2, this could decrease PRPP synthesis and thereby enhance PurR’s DNA binding activity.

Based on previous studies, there is no clear evidence in *B. subtilis* that the catalytic residues of RNase J2 are directly involved in RNA degradation *in vivo*. Our data suggest that in contrast to RNase J2 deletion, mutagenesis of its catalytic site (H76A) has no impact on RNA stability, at least on tested targets. Additionally, the pellicle of this mutant exhibits a slightly increased wrinkling compared to the wild-type strain, though this phenotype is far less pronounced than in the *ΔrnjB* strain. A structural impact of the H76A mutation on RNase J2 cannot be ruled out, which may subtly influence RNase J1 activity on specific substrates. However, no such effect was observed on the degradation of the tested targets. Notably, an equivalent mutation of the catalytic site in *S. aureus* did not result in the narrow-temperature-window-phenotype observed the *ΔrnjB* mutant strain (16). Furthermore, replacing the catalytic site of RNase J2 with that of RNase J1 entirely eliminates its residual 5’-exoribonuclease activity (11), suggesting that RNase J2 does not directly participate in RNA degradation *in vivo*. Although we cannot completely exclude a more direct role of RNase J2 under specific growth conditions and/or on particular RNA targets, our data leads us to propose that RNase J2 acts more as a specificity factor for RNase J1 than as an active ribonuclease. This association between an active enzyme and an inactive paralogous protein is found among ß-lactamase proteins in all domains of life.

In humans, pre-mRNAs are cleaved at the 3’-end by the U7 small nuclear ribonucleoprotein (snRNP), an RNA-protein complex that contained the active ribonuclease CPSF73 that interacts with an inactive paralogous protein CPSF100. Another example is the long-form RNase Z protein (Trz1), involved in tRNA 3’-processing eukaryotes, and which consists of a fusion of an active and an inactive ß-lactamase domain in a single protein (35).

Several non-exclusive hypotheses can be proposed for the role of RNase J2 in stimulating RNase J1 activity. 1) RNase J2 might serve as a stabilizer of RNase J1, as has been proposed in *S. aureus* (16). In this case, it might be expected that the destabilization of RNase J1 upon RNase J2 deletion in *B. subtilis* would strongly impact growth, as observed upon RNase J1 deletion. However, the *rnjB* deletion does not strongly impact growth in *B. subtilis* and actually results in a slight increase in RNase J1 levels (36). 2) RNase J2 could also help RNase J1 to degrade particular RNA targets by recognition of specific RNA sequences, structures or 5’ extremities and/or it could modify RNase J1 structure and activity to degrade these substrates more efficiently. These two hypotheses are consistent with our observation that mutating a single residue in J2 that contacts RNase J1 (R544) abolishes RNase J2’s contribution to RNA degradation and yields a biofilm phenotype indistinguishable from that of the *ΔrnjB* mutant. R544 resides in the C-terminal region of RNase J2, but it lies adjacent to the RNA-entry channel of RNase J1. A structural alteration in this area could impair RNase J2’s ability to deliver its specific substrates to the catalytic pocket of RNase J1, thereby compromising targeted RNA turnover. Interestingly, this residue was previously proposed to act as a switch for the positioning of the C-terminus (31). The mirror mutation of residue D449 in RNase J2, also predicted to form a salt-bridge with RNase J1 R544 and located near the predicted RNA-entry of RNase J2 (Fig. 8A) only slightly impact the degradation of RNase J2 targets such as the *dacA* mRNA, suggesting that this interaction is not necessary for an efficient RNA degradation. 3) RNase J2 could also modify RNase J1’s subcellular localization and hence its ability to find its targets. 4) Finally, RNase J2 could increase the number of active RNase J1 molecules without the need to increase its synthesis. In other words, at a comparable level of RNase J1 expression, heterodimerization with RNase J2 could increase the proportion of active dimers relative to RNase J1 homodimers, thereby potentially enhancing RNase J1 access to specific cellular targets. Consistent with this hypothesis, deletion of RNase J2 in *S. aureus* can be partially complemented by overexpression of RNase J1 or its catalytic mutant, indicating that dimer formation plays a critical role in maintaining catalytic activity (16). However, the fact that complementation is only partial also suggests that RNase J2 may contribute additional substrate specificity. The identification of the molecular mechanisms underlying the stimulation of RNase J1 by RNase J2 are under investigation.

## Materials & Methods

### Strains

All strains used in this study are derivative from *Bacillus subtilis* NCIB3610 comI-Q12L. Deletions of RNase J1 and RNase J2 was performed by transforming this background with chromosomal DNA from laboratory strains *B. subtilis* 168 CCB651 *(ΔrnjA::kan*) and CCB1561 *(ΔrnjB::spc*), respectively.

The complemented strain for RNase J2 was made by amplifying *rnjB* gene using primers CC3134/CC3135. The catalytic mutant of RNase J2 was generated by cloning a mutated version of the *rnjB* gene (H76A), amplified from pl513 carrying the H76A point mutation using primers CC3134/CC3135. The interacting mutants of RNase J2 (D449K and R544E) were amplified using the following oligo pairs: CC3134/CC3559 and CC3558/CC3135 for the D449K mutant, and CC3134/CC3561 and CC3560/CC3135 for R544E mutant. Amplifications were performed from pl1049, which carries the wild-type version of *rnjB*.

Cloning was performed using NEBuilder HiFi DNA assembly, following the manufacturer’s instructions (New England biolabs) into the BamHI site of the pDG1662 plasmid (pl315). The resulting plasmids (pl1049, pl988, pl1070 and pl1071) were then used to integrate the *rnjB* and its mutants at the *amyE* locus of the CCB1626 strain*(ΔrnjB::spc*).

The GFP-labeled strains were constructed by amplifying the *gfp* gene using oligo pairs CC2889/CC2422 and CC2890/CC572 from plasmides pl869 and pl899, respectively, and described in (37). An overlapping PCR was then performed using the oligo pairs CC2889/CC572. The resulting PCR fragment was cloned into pHM2 plasmid, previously digested with EcoRI and HindIII. The resulting plasmid (pl903) was used to integrate *gfp* at the *amyE* locus of the CCB1624 and CCB1626 strains, generating the CCB1805 and CCB1806 strains, respectively.

**Table S2.** Bacterial strains and plasmids.

| Table S2: Bacterial strains and plasmids |  |  |
| --- | --- | --- |
| Strain | Relevant genotype (drug resistance) | Source or reference |
| CCB1624 | NCIB3610 coml-Q12L (WT) | Konkol et al., 2013 |
| CCB1626 | NCIB3610 coml-Q12L *rnjB::spc* (*spc*<sup>R</sup>) | This study |
| CCB1719 | NCIB3610 coml-Q12L *rnjA::kan* (*kan*<sup>R</sup>) | This study |
| CCB1726 | NCIB3610 coml-Q12L *rnjB::spc amyE::pDG1662-rnjB* (H76A) (*Spc*<sup>R</sup> ; *Cm*<sup>R</sup>) | This study |
| CCB1728 | NCIB3610 coml-Q12L *rnjB::spc rnjA::kan* (*Spc*<sup>R</sup> ; *kan*<sup>R</sup>) | This study |
| CCB1805 | NCIB3610 coml-Q12L *amyE::Pspac<sup>con</sup>GFP-cm* (from pl903) | This study |
| CCB1806 | NCIB3610 coml-Q12L *amyE::Pspac<sup>con</sup>GFP-cm* (from pl903) | This study |
| CCB1904 | NCIB3610 coml-Q12L *rnjB::spc amyE::pDG1662-rnjB-CHis* (*Spc*<sup>R</sup> ; *Cm*<sup>R</sup>) | This study |
| CCB1964 | NCIB3610 coml-Q12L *rnjB::spc amyE::pDG1662-rnjB* (R544E) (*Spc*<sup>R</sup> ; *Cm*<sup>R</sup>) | This study |
| CCB1963 | NCIB3610 coml-Q12L *rnjB::spc amyE::pDG1662-rnjB* (D449K) (*Spc*<sup>R</sup> ; *Cm*<sup>R</sup>) | This study |
| Plasmid | Drug resistance (*B. subtilis* / *E. coli*) | Source or reference |
| pl315 | pDG1662 | Guérout-Fleury et al., 1996 |
| pl513 | pET28-*rnjB*(H76A) *rnjA-CHis* |  |
| pl903 | pHM2-Pspac<sup>con</sup>-GFP; used to construct CCB1805 and CCB1806 (*Ap*<sup>R</sup>) | This study |
| pl988 | pDG1662-*rnjB* (H76A) ; used to construct CCB1726 (*Cm*<sup>R</sup>/*Ap*<sup>R</sup>) | This study |
| pl1049 | pDG1662-*rnjB-CHis*; used to construct CCB1904 (*Cm*<sup>R</sup>/*Ap*<sup>R</sup>) | This study |
| pl1070 | pDG1662-*rnjB-CHis* (D449K) ; used to construct CCB1963 (*Cm*<sup>R</sup>/*Ap*<sup>R</sup>) | This study |
| pl1071 | pDG1662-*rnjB-CHis* (R544E) ; used to construct CCB1964 (*Cm*<sup>R</sup>/*Ap*<sup>R</sup>) | This study |

### Oligonucleotides

**Table S3.**
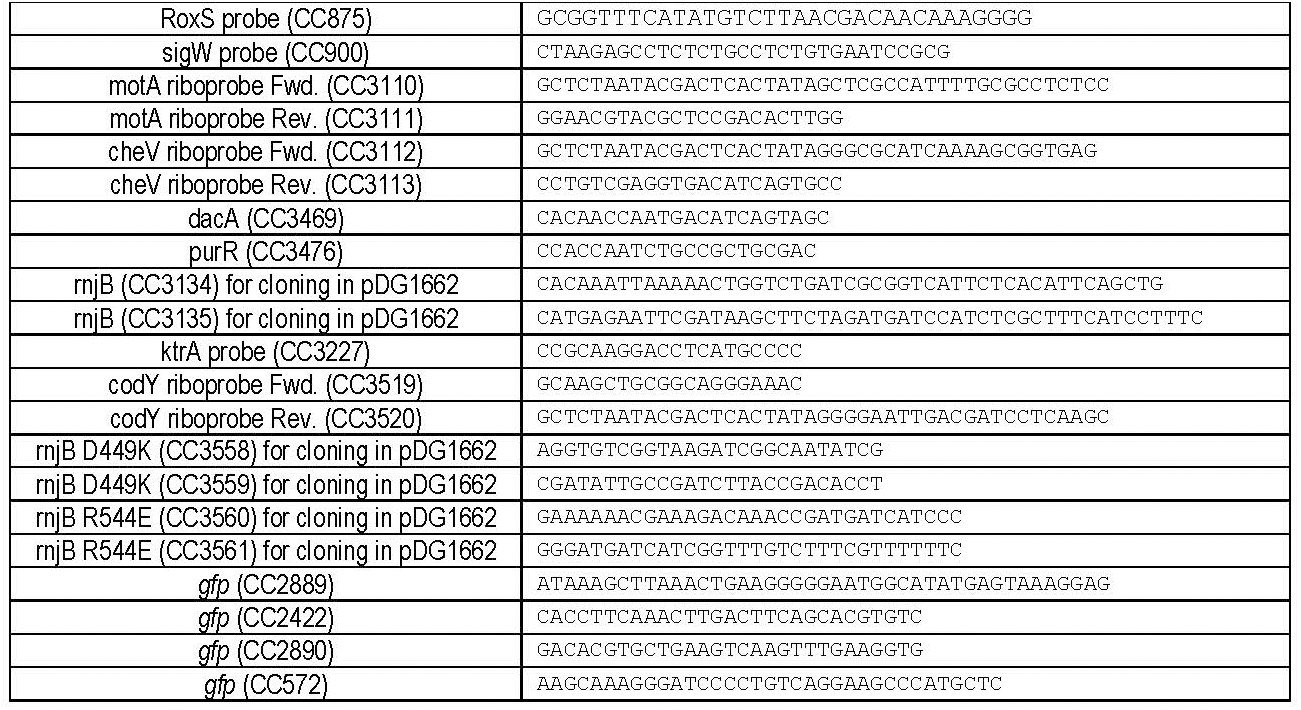
Oligonucleotides (sequences are 5’→3’)

### Biofilm growth

Two different biofilm models were used to characterize the phenotypes of our mutant and parental strains in this study: the macrocolony and the pellicle. For sample preparation of both biofilm models, an overnight culture grown in 2YT medium was diluted to an OD₆₀₀ of 0.05 in 2YTGM medium (composed of 2YT supplemented with 0.1 mM MnSO₄ and 1% glycerol). Cultures were incubated at 37 °C with agitation at 180 rpm until reaching the mid-exponential growth phase. Bacterial suspensions were then diluted to an OD₆₀₀ of 0.01 and used for macrocolony inoculation. A 2 µL drop of the bacterial suspension at mid-exponential phase was deposited onto 2YTGM agar plates containing 1.5% agar. Each well of a 12-well polystyrene plate (untreated, SARSTEDT) was filled with 2 mL of medium for macrocolony culture. Images of macrocolonies were acquired every hour using the automated plate imaging system Reshape, which integrates AI-based analysis to extract macrocolony area and growth rate. Data were analyzed using the Baranyi and Roberts growth model coupled with a custom Python script. The same protocol was used for swarming assays, except that the 2YTGM medium contained 1% agar. For pellicle formation, each well of a 24-well polystyrene plate (untreated, SARSTEDT) was filled with 1.5 mL of bacterial suspension at mid-exponential phase diluted to an OD₆₀₀ of 0.01. For both biofilm models, plates were sealed with parafilm to ensure homogeneous oxygenation across the wells.

### Confocal laser sacnning Microscopy

For confocal microscopy imaging, pellicle growth was performed following the protocol described above, using polystyrene 96-well microtiter plates with a μClear base (Greiner Bio-One, France), which are compatible with microscopy. Image acquisition was carried out in a time-resolved (kinetic) manner, from the onset of biofilm formation (t₀) up to 18 h. The strains used were CCB1805 and CCB1806, corresponding to the mutant and parental strains, respectively, both constitutively expressing GFP, allowing fluorescence measurements without the need for additional staining. Confocal imaging was performed using a Leica SP8 AOBS inverted high-content screening confocal laser scanning microscope (HCS-CLSM, Leica Microsystems, Germany). All acquisitions were conducted at the MIMA2 platform (https://www6.jouy.inra.fr/mima2_eng/). Z-stack images were acquired across the entire height of the biofilm, from the submerged layer at the bottom of the well to the pellicle located at the air–liquid interface. Images were collected at a resolution of 512 × 512 pixels (147.62 × 147.62 µm) using a 10× dry objective lens (numerical aperture = 0.30). Acquisition was performed at a scanning frequency of 400 Hz.

GFP fluorescence was excited using an argon laser at 488 nm, and emission was detected using a hybrid photomultiplier tube detector (HyD PMT, Leica Microsystems, Germany) in the range of 500–550 nm.

### Stastistical analysis and software

Biofilm development was quantified using GraphPad Prism 9 (GraphPad Software, version 9.0.0, San Diego, CA, USA) for all data processing and graphical representations. For biofilm experiments, data distribution was first assessed using the Shapiro–Wilk test to evaluate normality. Normally distributed data were compared between groups using a parametric Student’s t-test, while non-normally distributed data were analyzed using a non-parametric Mann–Whitney test.

### RIF-Seq

#### Sampling

*B. subtilis* NCIB3610 comI-Q12L (CCB1624) and NCIB3610 comI-Q12L *rnjB*:*:spc* (CCB1626) were cultivated in 2YT medium at 37°C until mid-exponential phase (O.D._600_ = 0.6). A sample of 10 ml from each culture was collected prior to the addition of rifampicin (at a final concentration of 150 µg/mL). Additional samples were taken at 1, 2, 5 and 10 min following rifampicin addition.

Cultures were done in triplicate from different precultures of each strain (biological replicates).

RNA were extracted by phenol-chloroform.

#### Sequencing, Alignment and normalization

RNA samples were sequenced in paired-end (2×150 bp) at the Institut du Cerveau, Hôpital Pitié-Salpêtrière, Paris with NextSeq2000 sequencing system. They performed rRNA depletion, trimming and demultiplexing of the data.

Paired-end reads are mapped on the NCIB3610 reference genome (GCF_002055965.1) using bowtie2 v2.5.0 (--very-sensitive-local). Quantification is performed using featureCounts v2.0.3 using annotation composed by the reference annotation file GCF_002055965.1 and homologous “segments” and “regulatory RNA” features identified by blast from strain 168 (subtiwiki, GCF_000009045.1), first normalized by gene length (cpk) to allow intra sample comparisons. A second normalization is then performed on three known stable transcripts (*ssrA*, *hag* and *rnpB,* see Fig. S7) to correct the massive overall transcript destabilization upon rifampicin treatment. Finally, scalling factor (ratio of median) between time 0 are applied to perform inter strain abundance comparison at t0. Genes with less than 10 counts in all samples are discarded.

#### RNA decay

Relative logarithmic RNA abundances were fitted using a weighted linear regression model with squared weights. To assess the quality of the fitting, a Mean Absolute Error (MAE) was calculated for the linear regression. For Half-life comparisons, only values with a MAE ≤ 0.6 in both the WT and the *rnjB* mutant strains were considered. This threshold was selected because *purR* has a MAE of 0.6 in the WT, and its calculated half-life remains consistent with that determined by Northern blot analysis (Fig. 5). Half-lives were considered significantly affected when the fold-change (FC) was ≥ 1.5 and the p-value was ≤ 0.05.

### Northern blot analyses

RNA was isolated from mid-log phase *B. subtilis* cells –grown in rich medium (2XYT) by the RNAsnap method, as described previously. Typically, 5 µg RNA was run on 1% agarose in 1X TBE buffer and transferred to Hybond-N+ membranes (Cytiva) by capillarity. Hybridization was performed using either 5’-labeled oligonucleotides labeled with [g-^32^P]-ATP and T4 polynucleotide kinase, following manufacturer’s instructions or a riboprobe transcribed from a PCR-amplified DNA template containing a T7 promoter and labeled during T7 transcription with [a-^32^P]-UTP. Hybridization was carried out at 42°C (for oligonucleotide probes) or 68°C (for riboprobes) for a minimum of 4 h in UltraHyb hybridization buffer (Ambion). Membranes were washed twice in 2X SSC/0.1% SDS (once rapidly at room temperature and once for 10 min at 42°C) and then three times for 10 min in 0.2X SSC/0.1% SDS at room temperature.

### Western blot analyses

Proteins were isolated from mid-log phase *B. subtilis* cells –grown in rich medium (2XYT). Lysis was performed by sonication in TE-NaCl lysis buffer (10 mM Tris-HCl pH 8; 1 mM EDTA; 0.1 M NaCl; 1 MiniComplete EDTA-free protease inhibitor tablet) and protein concentration assessed by Bradford assay (Bio-Rad Protein Assay kit). Typically, 20 µg of proteins were loaded on an 8% acrymalide-bisacrylamide gel and run in SDS-Tris-Glycine running buffer before electrophoretic transfer to nitrocellulose membrane in transfer buffer (10% ethanol; 10 mM Tris base; 192 mM glycine). The membrane was then blocked in PBST (0.1% Tween-20) containing 5% milk and incubated with anti-Flag antibody (1:1000) in PBST followed by incubation with HRP-conjugated anti-rabbit secondary antibody (1:10,000). Detection was carried out by chemiluminescence using the Covalight kit (Covalab) on a Chemidoc system (BioRad).

## DATA AVAILABILITY

Raw RNA-seq data (FASTQ files) will be deposited in the Gene Expression Omnibus (GEO) database under accession number [XXX].

## ACKNOWLEDGMENTS

We acknowledge the sequencing and bioinformatics expertise of A.M of the IBPC functional genomics facility. We are grateful to Daniel Kearns for providing the *Bacillus subtilis NCIB3610* strain and to Elena and Vladimir Bidnenko for providing the *B. subtilis* strain CYR9.

## AUTHOR CONTRIBUTIONS

N.C., M.G., T.K. performed experiments; A.M, H.L. and N.C. made the bioinformatic analysis S.D., designed the study and wrote the manuscript with the help from the other authors.

## FUNDING

This work was supported by the French Agence Nationale de la Recherche (ANR-24-CE12-5113 to S.D.) and by the LABEX program Dynamo (S.D.). N.C is the recipient of a PhD fellowship from the LABEX program Dynamo.

## LICENSE

CC-BY

## CONFLICT OF INTEREST

The authors declare no competing interests.

## SUPPLEMENTAL INFORMATION

### Supplementary Figures

**Supplementary Figure 1.**
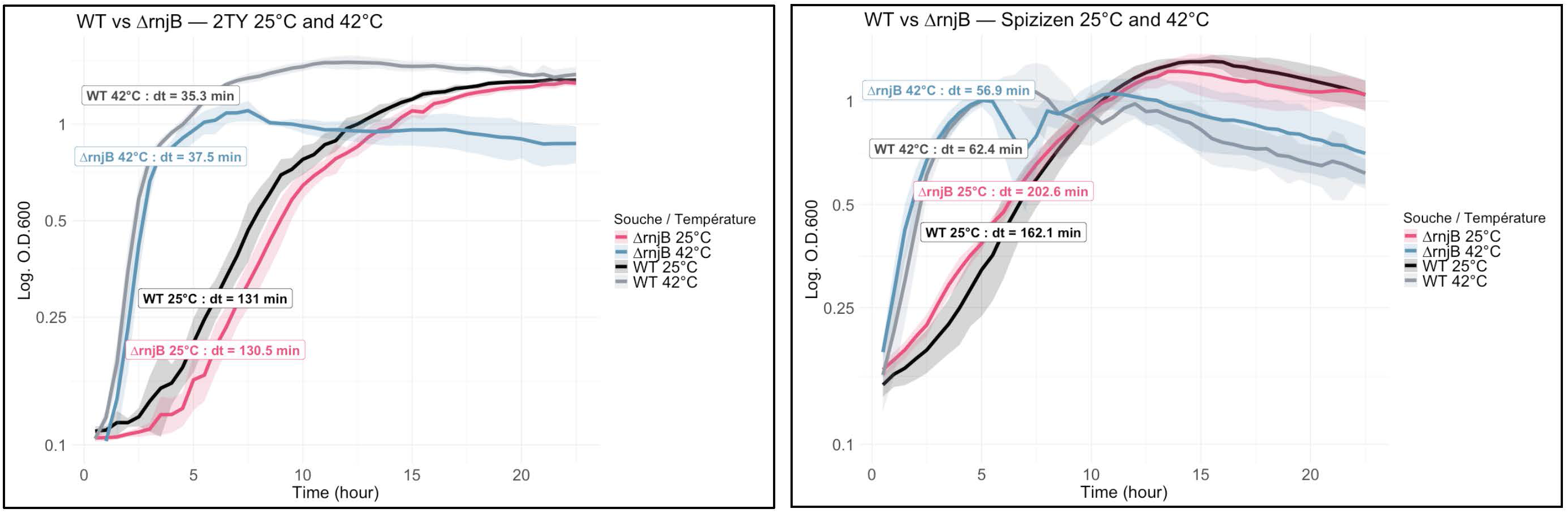
Growth curves of the WT and ΔrnjB strains Strains were innoculated into 96-well plates from an overnight culture adjusted to O.DD.600=0.05 in 2YT medium (A) and in Spizizen minimal medium (B). Growth was monitored at 25°C and 42°C, with O.D.600 readings taken every 30 min. dt: doubling time

**Supplementary Figure 2.**
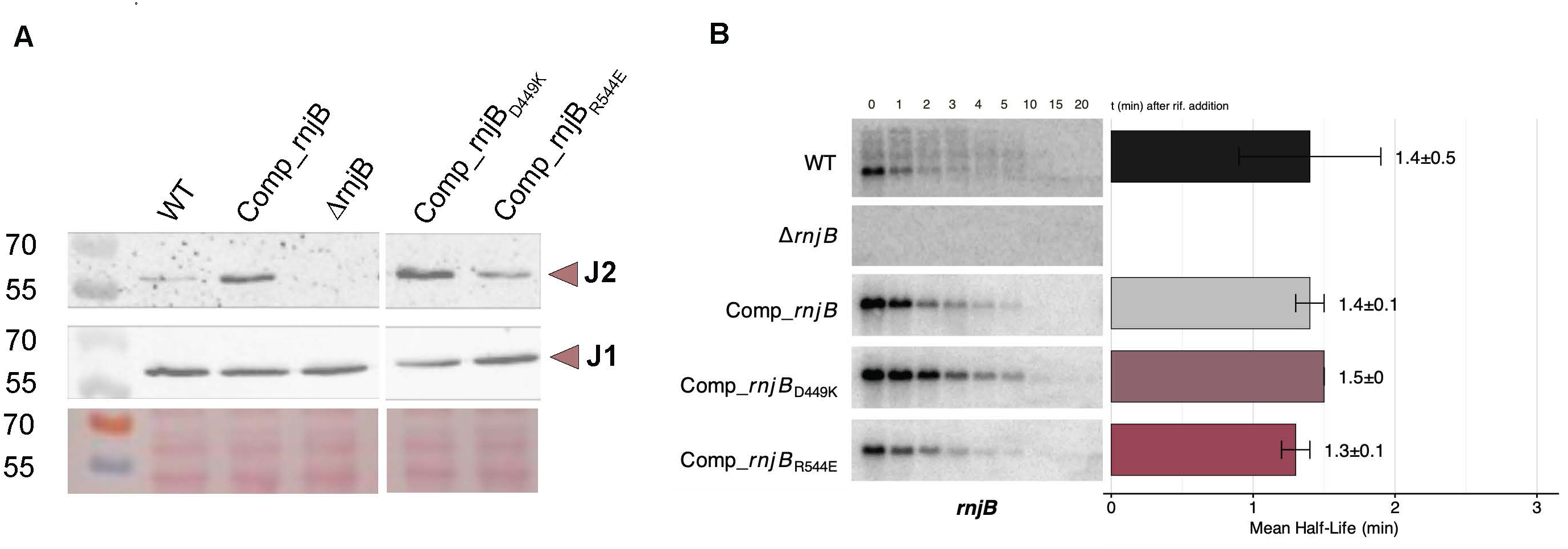
RNase J2 protein and mRNA levels are similar in wild-type (WT) and ΔrnjB complemented strains. (A) Western blot analysis using RNase J1 and J2 antibodies. Total protein samples were extracted from cells grown in 2YT medium at mid-exponential phase in the following strains: WT, ΔrnjB and ΔrnjB strains complemented strains at the amyE locus with rnjB-CHis WT, D449K, or R544E. Ponceau S staining below each lane serves as a loading control to confirm equal protein loading. (B) Northern blot analysis of total RNA from WT, ΔrnjB and ΔrnjB complemented strains (WT, D449K, R544E), probed with a rnjB radiolabeled probe. All samples for Northern blots were collected at mid-exponential phase in 2YT medium, before (0) and after rifampicin treatment (1 to 20 min time points).

**Supplementary Figure 3.**
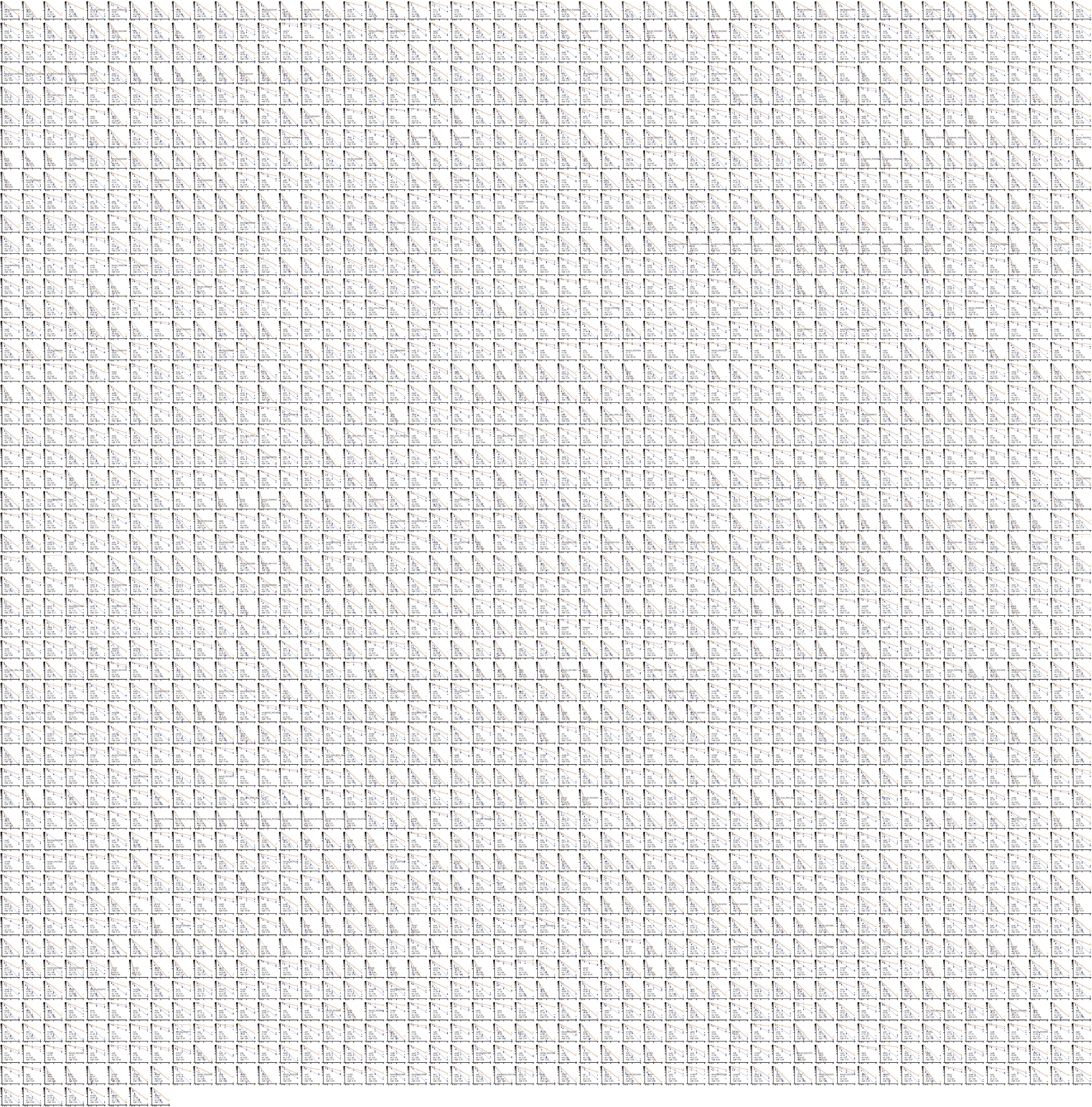
Comparison of RNA half-lives on a logarithmic scale between wild-type (WT) and ΔrnjB strains. Relative logarithmic RNA abundances were fitted using a weighted linear regression model with squared weights. Individual data points represent biological replicates for WT (blue) and ΔrnjB (orange). The half-life for each strain (wt and ΔrnjB) and the fold-change (x) are indicated on the graph for each RNA.

**Supplementary Figure 4.**
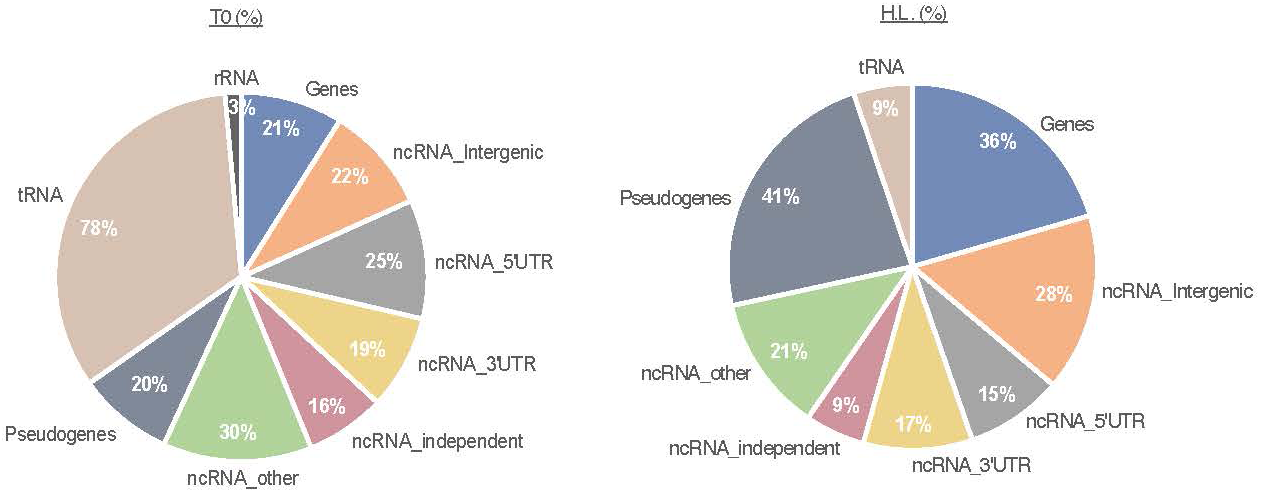
Impact of rnjB deletion on RNA features. Percentage of RNAs for each feature (as described in Nicolas et al., 2012) significantly affected in the ΔrnjB compared to the WT strain at T0 (T0) or in their half-lives (HL) (|log2FC|≥ 0,585, p-value ≥ 0.05 for both T0 and HL, additionnaly HL changes were filtered using MAE ≤ 0.6).

**Supplementary Figure 5.**
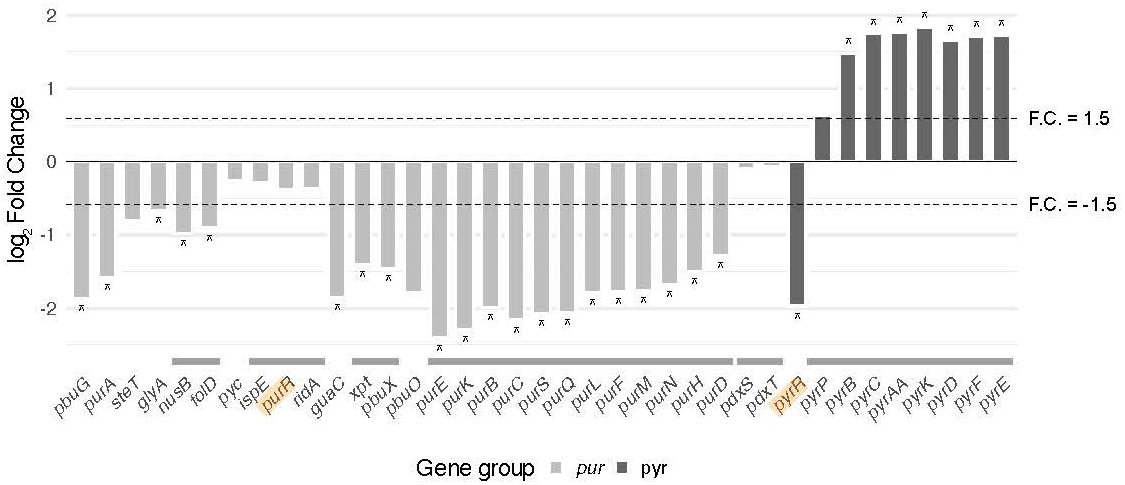
Differential gene expression within the PurR and PyrR regulons in WT and ΔrnjB strains at steady-state (T0). Operons are represented with a grey line above the gene name. Stars indicate significant differences (|log2FC_t0 |≥ 0.585 and p-value ≤ 0.05). Dashed lines indicate |log2FC_t0 |≥ 0.585. Genes encoding transcriptional regulators PurR and PyrR are highlighted in yellow.

**Supplementary Figure 6.**
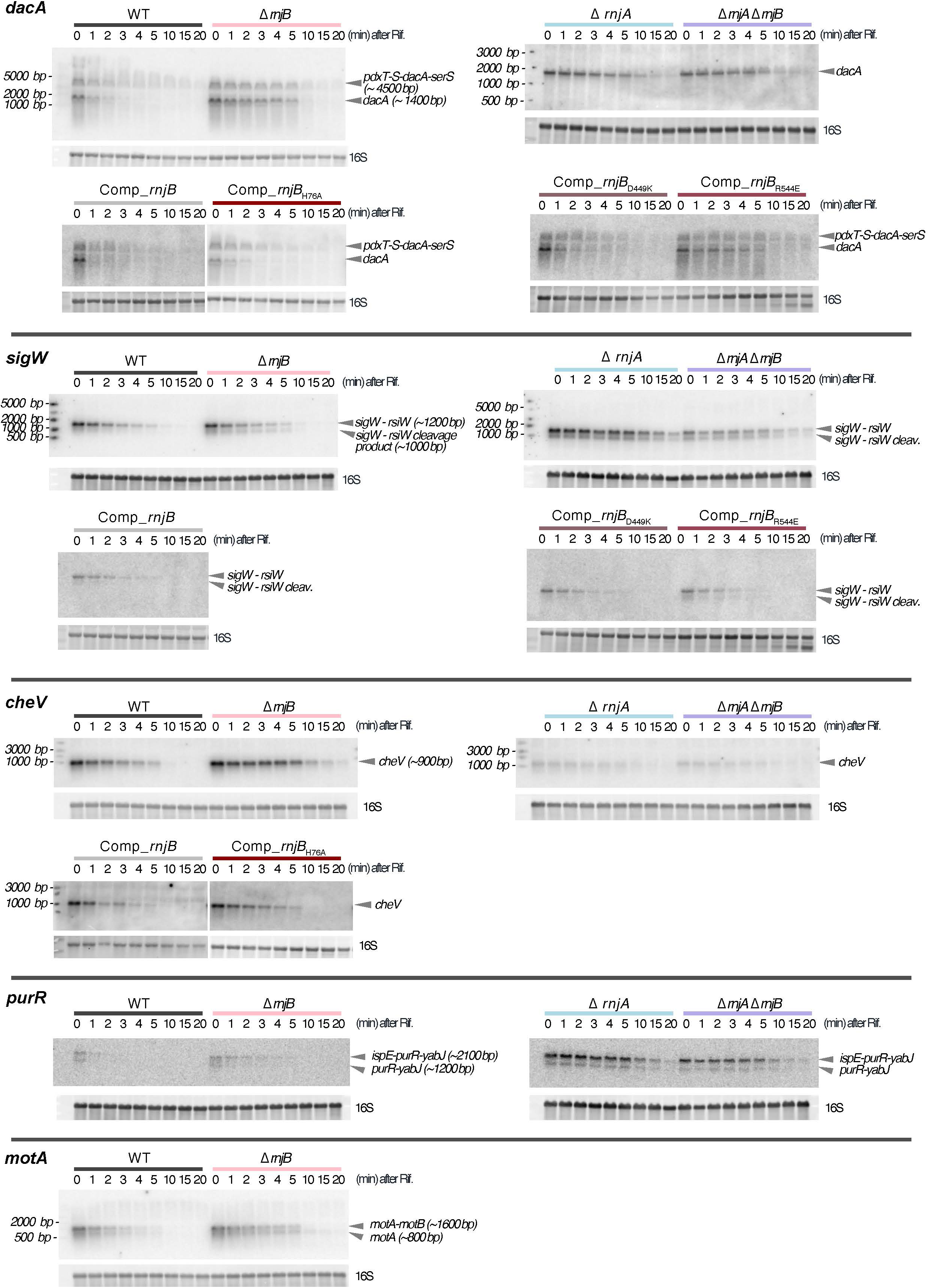
Representative Northern blots used for half-life determination. Cells were grown to mid-exponential phase (O.D._600_=0.7). Total RNA was extracted from WT, ΔrnjA, ΔrnjB, and complemented strains expressing either wild-type rnjB (Comp_rnjB), the catalytic mutant (Comp_rnjB_H76A_) or interaction mutants (Comp_rnjB_D449K_ or Comp_rnjB_R544E_). Samples were collected before (0) and after rifampicin addition at the indicated time points (1’, 2’, 3, 4’, 5’, 10’, 15’, 20’). Northern blots were hybridized with radiolabelled probes.

**Supplementary Figure 7.**
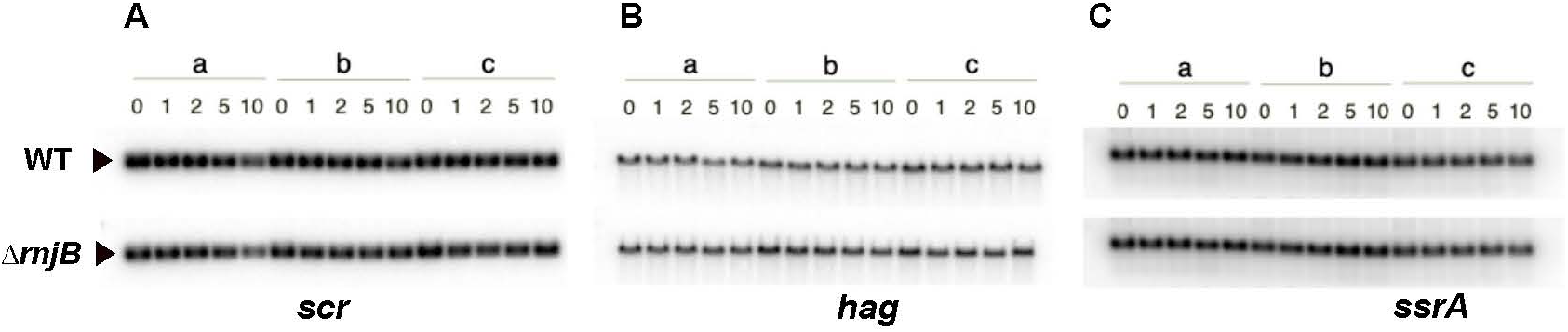
Control RNAs used for normalisation. Northern blot analysis of the total RNA sent for sequencing. The RNA were extracted from three biological replicates (a, b and c) of WT and ΔrnjB strain grown at mid-exponential phase in 2YT before (0) and after rifampicin treatment (1, 2, 5 and 10 min time points) to block transcription. The membranes were probed for (A) scr (B) hag and (C) ssrA using radiolabelled probes.

## Notes

### Competing Interest Statement

The authors have declared no competing interest.

### Summary of Updates

The author list has been revised.

